# Sequencing depth overcomes extraction bias: repurposing human WGS data for salivary microbiome profiling

**DOI:** 10.64898/2026.03.27.714786

**Authors:** Lourdes Velo-Suárez, Anthony F. Herzig, Ozvan Bocher, Gaëlle Le Folgoc, Liana Le Roux, Christelle Delmas, Marie Zins, Jean-François Deleuze, Geneviève Héry-Arnaud, Emmanuelle Génin

## Abstract

Large-scale human genomic projects have generated whole-genome sequencing (WGS) data from hundreds of thousands of individuals, primarily to study host genetic variation. When saliva is the DNA source, the resulting datasets also contain microbial reads that are routinely discarded. Here, we investigate whether these host-centric WGS workflows can yield reliable microbiome profiles, effectively doubling the research value of existing data without additional sampling. We compared non-human reads from 39 deeply sequenced saliva samples from the GAZEL cohort (miG dataset; median ∼43 million reads/sample) with 14 samples processed with microbiome-optimized extraction (ASAL; median ∼4.3 million reads/sample), using two complementary classifiers: meteor, a coverage-based mapper against a curated saliva-specific database, and sylph, a k-mer classifier against the Genome Taxonomy Database (GTDB). Despite the absence of microbial lysis optimization, miG samples showed up to 3-fold higher species richness, ∼10-fold greater sequencing depth, and significantly lower inter-sample variability (PERMANOVA R² = 0.10, p = 0.001; BETADISPER p = 0.0036). Rarefaction to 10⁶ reads eliminated most compositional differences, demonstrating that sequencing depth is the primary driver of community stability. Only ∼2% of detected taxa (12 of 592) showed extraction-related differences. The two classifiers exhibited fundamentally different depth-sensitivity profiles, with sylph retaining systematic detection asymmetries even after depth normalization, highlighting that classifier choice introduces biases that affect cross-study comparisons. These results show that biobank WGS data from saliva can be repurposed for robust, population-scale oral microbiome analyses, enabling simultaneous investigation of host genomic variation and the microbiome from the same archived samples.

**Importance:** Saliva-based whole-genome sequencing datasets generated across various cohorts to study human genetics contain non-human reads that are routinely discarded, thereby overlooking valuable microbial information. We show that these reads are sufficient to reconstruct robust oral microbiome profiles — without any additional sampling or laboratory work. This finding unlocks a vast archive of existing genomic data for retrospective microbiome research, enabling population-scale studies of oral microbial diversity, host–microbiome interactions, and disease associations at minimal additional cost. We further demonstrate that the choice of taxonomic classifier introduces systematic, depth-dependent biases that persist even after normalization, a practical consideration for any cross-cohort or multi-platform microbiome study.

## Introduction

Large-scale human genomic biobanks, including the UK Biobank (1), FinnGen (2), Fin-HIT (3), SPARK (4), and All of US (5), have collectively generated whole-genome sequencing (WGS) data from millions of individuals, primarily to map host genetic variation and disease associations. When saliva serves as the DNA source, each sequenced sample contains not only host reads but also a microbial fraction that is routinely removed and discarded during variant-calling pipelines (6). This constitutes an enormous and largely untapped archive of microbiome data: retrospective microbiome profiling from existing biobanked samples would require no additional collection, no renewed consent for sampling, and no additional laboratory processing, while enabling the integration of host genomic and microbial data within the same individuals. The key question is whether these host-optimized workflows can produce reliable microbiome profiles.

The salivary microbiome, a dynamic and diverse microbial ecosystem within the oral cavity, plays a vital role in both oral and systemic health. Saliva supports essential physiological processes, including digestion, mucosal immunity, and microbial homeostasis, and offers a non-invasive window into host–microbe interactions (7–9). Increasing evidence links salivary microbial signatures to diseases and immune modulation, positioning saliva as a valuable biospecimen for population-scale health surveillance and microbiome research (10, 11).

Salivary microbial community profiling has traditionally relied on 16S rRNA gene profiling (12, 13), which involves targeted amplification of bacterial ribosomal genes after DNA extraction and yields primarily taxonomic information on community composition. In contrast, whole-genome sequencing (WGS) does not rely on targeted PCR amplification; it directly sequences all extracted DNA, capturing both host and microbial reads. This untargeted approach enables simultaneous taxonomic profiling and functional characterization of the microbiome. In practice, the shift from 16S rRNA profiling to WGS in microbiome analysis remains limited by cost and regulation. Because saliva contains about 90% human DNA(14), WGS is costly, and host-derived sequences impose legal and ethical constraints requiring authorization and secure data handling. As a result, salivary microbiome studies have historically relied on 16S rRNA or, recently, on shallow shotgun sequencing, which remain more affordable alternatives.

In WGS-based biobank settings, extraction protocols are typically optimized for human genomic yield, often at the expense of microbial recovery. This can bias taxonomic profiles, particularly by underrepresenting key or structurally resilient taxa (15–17). Prior studies have shown that extraction methods significantly shape microbial richness and composition, especially for low-abundance or hard-to-lyse microorganisms(18–21). Within the oral microbiome, comparative studies further indicate that both DNA extraction and collection methods can influence microbial profiles, although salivary communities remain relatively robust across methodological variations (22, 23). Moreover, the trade-off between microbial lysis efficiency and sequencing depth is poorly defined. While specialized protocols improve microbial recovery, deep sequencing may mitigate extraction limitations by capturing robust microbial signals even from suboptimally processed samples (24, 25).

Classifiers fall broadly into three paradigms. K-mer or sketch-based methods, such as Kraken2/Bracken and sylph, classify reads by exact or near-exact subsequence matches against large reference databases, achieving high recall but at the cost of lower precision, particularly for low-abundance taxa. Marker-gene methods, such as MetaPhlAn4 and mOTUs, rely on clade-specific conserved genes, yielding higher precision but reduced recall. Coverage– or alignment-based approaches, such as meteor, infer taxon abundance from genome-wide read mapping against curated, community-specific databases, trading broad taxonomic coverage for depth-robustness within a defined ecological niche.

These precision-recall tradeoffs have been extensively characterized by community-led benchmarking initiatives (26), most comprehensively in successive rounds of the Critical Assessment of Metagenome Interpretation (CAMI), specifically CAMI I and II (27, 28), where marker-gene methods ranked best for abundance estimation precision while k-mer tools showed superior recall but higher false-positive rates at the species level. Critically, however, these benchmarks were conducted on datasets with uniform or controlled sequencing depth. Whether these well-characterized architectural differences are further compounded by order-of-magnitude depth disparities, as is inherent to biobank-scale WGS repurposing, where host-optimized and microbiome-optimized samples may coexist in the same study, has not been systematically investigated.

One recent example illustrating both the promise and the unresolved methodological questions of this approach is Kamitaki et al.’s study, which profiled oral microbiomes from 12,519 individuals in the SPARK cohort (4) using unmapped reads from saliva-based WGS. Their study demonstrated that repurposed microbial reads could be clustered with oral rather than gut communities and that associations between host polymorphisms and the principal axes of microbiome variation were detectable at the population scale. As their results may inspire the re-analysis of unmapped reads from other cohorts that have collected saliva, it becomes critical to establish whether host-optimized extraction protocols produce profiles of sufficient reliability for such analyses, and whether that conclusion holds across classifier paradigms with fundamentally different sensitivities.

In this study, we address three questions directly relevant to any researcher considering retrospective microbiome profiling from existing biobank data: (1) How do coverage-based and k-mer classifiers differ in taxon detection sensitivity as a function of sequencing depth? (2) Does rarefaction normalization equalize classifier behavior across depth-mismatched datasets? (3) Which, if any, taxa show genuine extraction-protocol biases independent of sequencing depth? By answering these questions, we provide both empirical validation and a practical decision framework for leveraging the vast archives of saliva-based WGS data already generated by large-scale human genomics initiatives.

## Results and Discussion

By comparing 39 deeply sequenced saliva samples processed with standard host-focused protocols (miG dataset) to 14 samples from an independent study extracted with microbiome-specific methods (ASAL dataset), we assess differences in taxonomic detection, community richness, and inter-sample variability—thereby testing the potential of high-depth, host-centric workflows for metagenomic microbiome research and integrated host–microbiome analyses. The overall analytical workflow is summarized in Figure 1.

**Figure 1.**
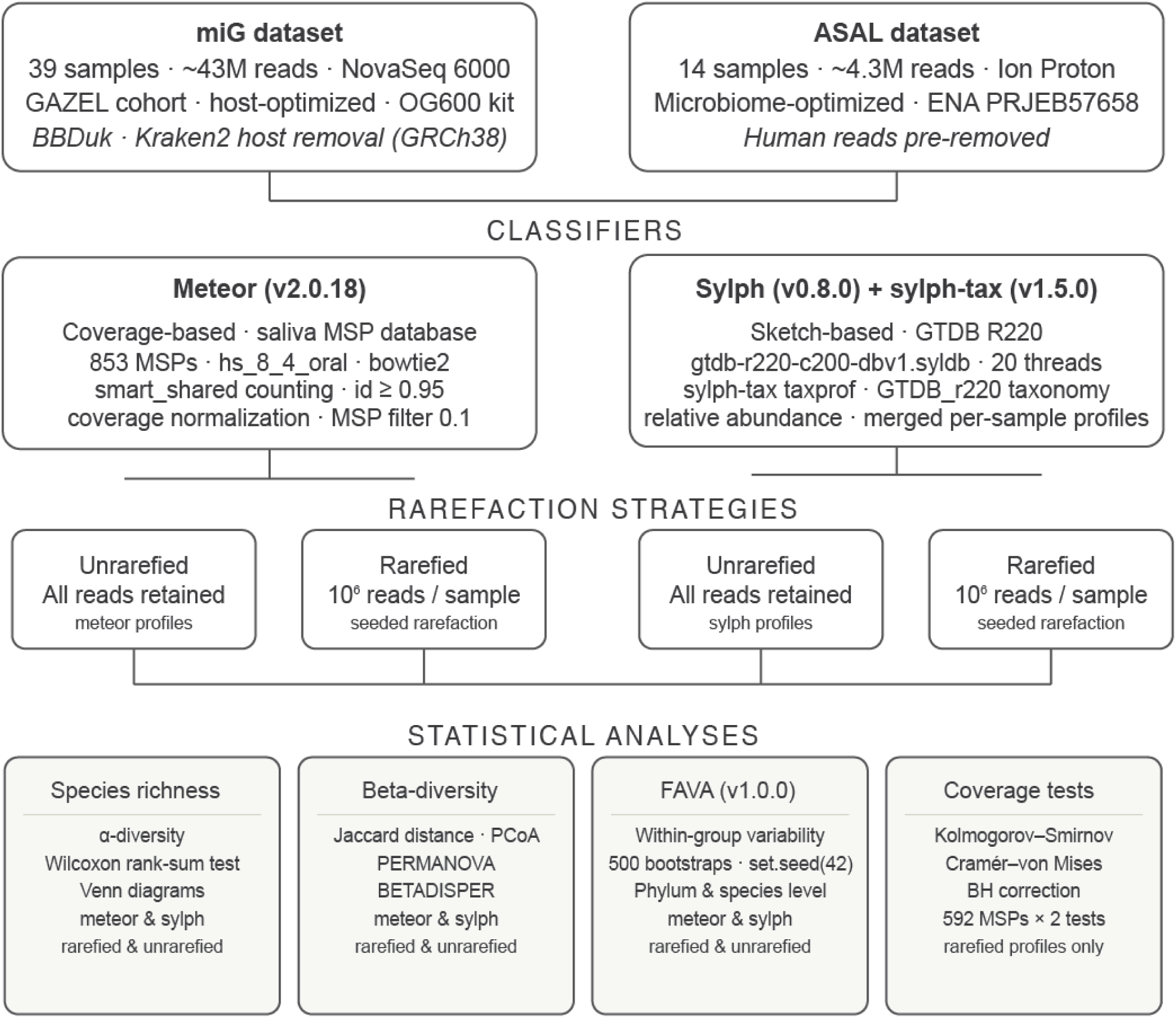
Bioinformatics workflow for comparative salivary microbiome profiling. Non-human reads from the miG dataset (39 saliva samples from the GAZEL cohort, sequenced at median ∼43 million reads/sample on an Illumina NovaSeq 6000) and the ASAL dataset (14 samples sequenced at median ∼4.3 million reads/sample on an Ion Proton platform; ENA accession PRJEB57658) were profiled in parallel using two complementary classifiers: meteor (v2.0.18), a coverage-based mapper against a curated saliva-specific database of 853 metagenomic species pangenomes (MSPs); and sylph (v0.8.0) with sylph-tax (v1.5.0), a sketch-based classifier against the Genome Taxonomy Database (GTDB release R220). Each classifier was applied to both rarefied (10⁶ reads/sample) and unrarefied data. Resulting profiles were evaluated through four statistical frameworks: species richness (α-diversity; Wilcoxon rank-sum test), microbial community structure (β-diversity; Jaccard distance, PCoA, PERMANOVA, BETADISPER), within-group compositional variability (FAVA v1.0.0; 500 bootstrap replicates, set.seed(42)), and per-MSP coverage distributions (Kolmogorov–Smirnov and Cramér–von Mises tests; Benjamini–Hochberg correction across 1,184 tests). BH, Benjamini–Hochberg; MSP, metagenomic species pangenome; GTDB, Genome Taxonomy Database; FAVA, F-statistic-based Analysis of Variability in Abundances.

### Sequencing depth differences between miG and ASAL datasets

The total number of quality-filtered reads varied between datasets, with the miG samples showing higher sequencing depth than the ASAL samples. The miG protocol yielded between 8.2 million and 196 million reads per sample (median approximately 43 million), while ASAL samples ranged from 1.5 million to 17 million reads (median approximately 4.3 million). Sequencing depth is a key determinant of taxonomic detection sensitivity and compositional stability, particularly in metagenomic studies where low-abundance taxa may be inconsistently captured across samples (25). The tenfold difference in sequencing depth between the miG and ASAL datasets provides essential context for interpreting differences in richness, variability metrics, and rarefaction effects. This depth advantage, a direct consequence of the deep sequencing required for rare-variant discovery in biobank studies, emerges here as an unexpected asset for microbiome recovery.

### Taxonomic detection and richness across profiling methods

We compared overall taxonomic detection and microbial community structure between the ASAL and miG datasets using two complementary profiling approaches: meteor and sylph. In unrarefied data, mean richness was significantly greater in miG for both meteor and sylph (Figure 2A and Figure 2B). After rarefaction to 10⁶ reads per sample, detection in meteor-profiled data became comparable between groups (Figure 2C), while miG samples still retained a clear advantage in sylph (Figure 2D). These results align with prior studies demonstrating that sequencing depth significantly influences taxon detection, particularly for low-abundance or difficult-to-lyse taxa (15, 18, 19).

**Figure 2.**
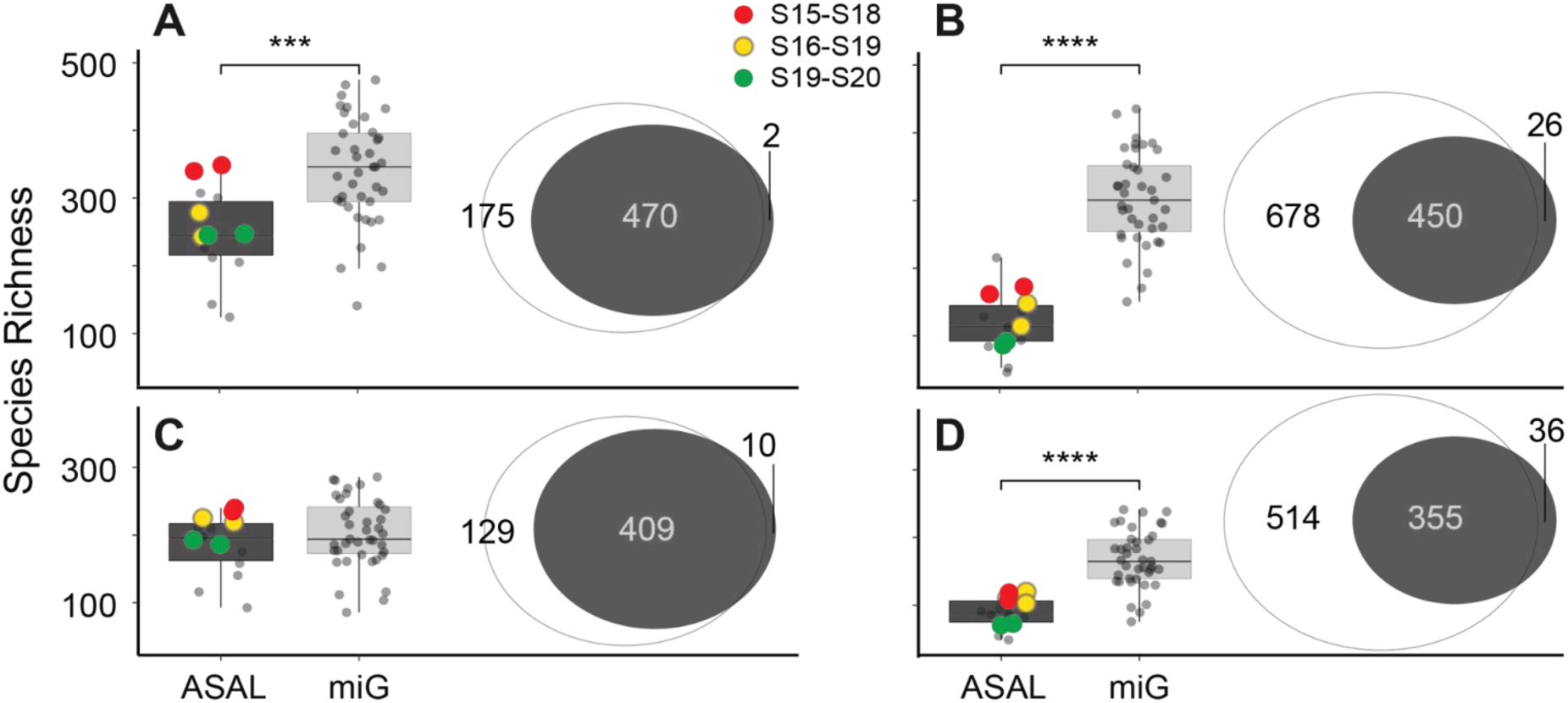
Species richness and taxon detection in ASAL and miG saliva samples. Profiles generated with meteor (left panels) and sylph (right panels) for unrarefied data (A, B) and rarefied data at 10⁶ reads (C, D). Venn diagrams show the number of shared and unique taxa between datasets. Colored points in the ASAL cohort represent pooled samples, with each color corresponding to a specific donor pair (red = S15–S18, yellow = S16–S19, green = S19–S20).

### Influence of database breadth and profiling strategy on taxon detection

Taxon detection is strongly shaped by both database breadth and profiling approach, with genome mapping (meteor) and k-mer classification (sylph) yielding distinct results. Across all conditions, per-sample richness was consistently lower with sylph than with meteor (Figure 2). This reduction reflects sylph’s reliance on strict k-mer thresholds across the broad GTDB reference, which restricts detection to high-confidence matches and therefore yields fewer taxa per individual sample. In contrast, meteor leverages a curated saliva-specific database and coverage-based detection, allowing for the more permissive recovery of low-abundance taxa and thus richer per-sample profiles. The average mapping rate to the meteor oral database was high in both groups (83.4% for miG and 82.0% for ASAL), indicating its suitability for salivary microbiota profiling. The Venn diagrams clarify the apparent discrepancy between methods: meteor produced largely overlapping profiles between ASAL and miG, with depth-related differences that diminished after rarefaction. Sylph, by contrast, yielded highly asymmetric outcomes: in unrarefied data, miG contributed more than 600 taxa not seen in ASAL, and even after rarefaction, it retained over 500 unique detections, whereas ASAL contributed only a few dozen. The surplus of taxa detected by sylph in miG reflects rare or very low-abundance species that appear sporadically across individuals: they accumulate when all samples are pooled together, but do not contribute consistently to per-sample richness. Taken together, these results highlight a key methodological contrast. Meteor maximizes per-sample richness and generates stable, comparable profiles between groups once sequencing depth is controlled, showing that deep sequencing in miG can largely compensate for its host-focused microbial extraction. Sylph, in turn, systematically reports lower per-sample richness but accumulates a larger number of unique taxa across samples in deeply sequenced datasets — sporadic, low-abundance detections that appear in individual miG samples but not consistently enough to raise per-sample richness. This makes sylph particularly sensitive to sequencing depth asymmetries, strongly penalizing shallow sequencing despite the microbiome-optimized extraction of ASAL. Rarefaction further illustrates this interplay: normalizing both datasets to 10⁶ reads attenuated differences in meteor, yielding broadly comparable community profiles, but only partially reduced the imbalance in sylph, where miG still retained hundreds of unique detections.

### Microbial community composition and beta-diversity patterns

Building on the observed differences in taxonomic richness (Figure 2), we next assessed microbial community composition using Principal Coordinates Analysis (PCoA) based on Jaccard distances (Figure 3). In the unrarefied meteor-derived profiles (Figure 3A), ASAL and miG samples formed distinct clusters (PERMANOVA R² = 0.102, p = 0.001) with significant differences in within-group dispersion (BETADISPER p = 0.0036). ASAL samples showed a broader spread in ordination space, whereas miG samples were more tightly clustered. This beta-diversity pattern is consistent with the alpha-diversity results from meteor (Figure 2A), in which miG samples exhibited higher per-sample richness and a larger set of unique taxa. Deep sequencing, therefore, stabilizes miG profiles across individuals, whereas the lower richness of ASAL results in higher compositional turnover and greater dispersion.

**Figure 3.**
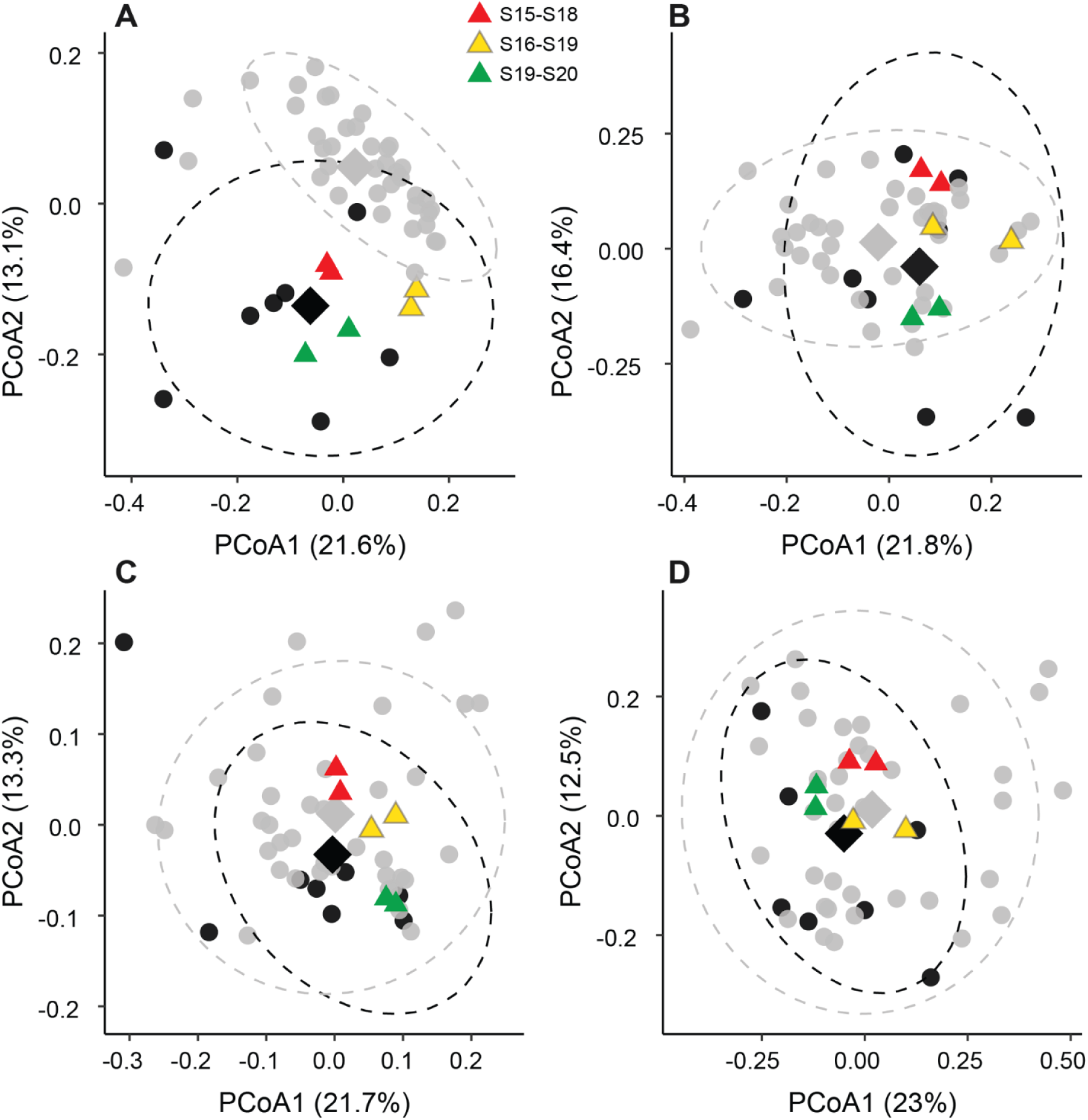
Microbial community structure in ASAL and miG saliva samples. Principal Coordinates Analysis (PCoA) plots based on Jaccard distances, profiled with **meteor** (left panels) and **sylph** (right panels). (A, B) Unrarefied data; (C, D) data rarefied to 10⁶ reads. Ellipses indicate 95% confidence intervals for each group. Group centroids are shown as rhombuses (black = ASAL, grey = miG), and pooled ASAL samples are marked with colored triangles. Pooled ASAL samples are indicated by colored triangles, with colors corresponding to the donor pairs included in each pool (red = S15–S18, yellow = S16–S19, green = S19–S20).

In contrast, the sylph-based profiles (Figure 3B) showed substantial overlap, with ASAL samples nested into the broader distribution of miG. Although separation remained statistically significant (PERMANOVA R² = 0.048, p = 0.013), dispersion differences disappeared (BETADISPER p = 0.12). This matches the sylph alpha-diversity results (Figure 2B), where miG again exhibited higher per-sample richness and hundreds of unique taxa, leading sylph to effectively position ASAL as a subset of the deeper miG profiles.

### Rarefaction effects on community structure

After rarefying all samples to 10⁶ reads, meteor and sylph showed different sensitivities to normalization. In meteor (Figure 3C), rarefaction removed many low-abundance taxa, disproportionately inflating richness in miG, leaving both datasets represented primarily by their shared core microbiota. This produced community profiles with small but statistically significant between-dataset differences (PERMANOVA R² = 0.045, p = 0.008) and no differences in dispersion (BETADISPER p = 0.589). In sylph (Figure 3D), ASAL profiles clustered within the broader miG distribution, consistent with the α-diversity patterns in Figure 2D, and statistical tests similarly indicated a small yet detectable separation (PERMANOVA R² = 0.046, p = 0.009; BETADISPER p = 0.681).

Together, these results show that rarefaction reduces between-dataset differences in both methods, leading to similarly small PERMANOVA effect sizes. However, the origin of this residual separation differs: in meteor, community structures nearly converge after depth normalization, whereas in sylph the remaining separation reflects its greater sensitivity to low-abundance taxa and the deeper sampling of the miG dataset. This classifier-dependent response to rarefaction is not an artefact of normalization but a structural property of sylph’s minimizer-based architecture against the broad GTDB reference: even at matched sequencing depth, sylph retains depth-acquired signal that coverage-based approaches like meteor do not.

These findings have a direct practical implication for researchers repurposing host-centric biobank WGS data. Rarefaction normalizes sequencing depth but does not equalize classifier behavior: the residual between-dataset asymmetry observed in sylph after rarefaction reflects its greater sensitivity to rare taxa and the architecture of its minimizer-based scoring against the broad GTDB reference, rather than true biological differences between cohorts. Community-led benchmarking has established that k-mer and sketch-based classifiers systematically report higher recall but lower precision than coverage– or marker-gene-based approaches, particularly at the species level (27, 28). Our results extend this observation to a biobank-realistic scenario — a host-contaminated matrix with a ten-fold depth mismatch — and show that this precision-recall tradeoff is not resolved by rarefaction. Classifier choice should therefore be treated as an independent analytical variable in cross-cohort microbiome studies using repurposed WGS data, not an interchangeable implementation detail.

### Minimal impact of the extraction protocol on community structure

Finally, in both diversity analyses, the pooled ASAL sample pairs were consistently highly similar, regardless of extraction protocol (manual or semi-automated). In Figure 2, the paired samples, shown as colored circles, display nearly identical richness values, while in Figure 3, the identical pairs—shown as colored triangles—cluster tightly together in ordination space. Together, these patterns indicate that the extraction method had minimal impact on overall community structure, supporting the conclusion that host-centric or semi-automated protocols can yield reliable salivary microbiome profiles. This is particularly relevant for large biobanks, where standardized, partially automated workflows are used for efficiency and where further optimization of microbial lysis would be impractical at scale.

### Within-group compositional variability assessed by FAVA

To further evaluate microbial community variability, we used FAVA (29), which quantifies within-group compositional consistency (Figure 4). Higher FAVA values indicate greater inter-individual compositional variability, and lower values indicate more stable microbial community structures within groups. Across taxonomic ranks and both profiling methods, rarefied miG, ASAL, and mixed datasets displayed similar FAVA distributions. Notably, the mixed group did not show higher variability than either cohort alone, indicating that combining samples processed under different extraction and sequencing conditions does not inflate inter-sample differences. FAVA values decreased from broad (phylum, class) to fine (genus, species) taxonomic levels, consistent with greater stability at higher ranks and increased sensitivity of species-level profiles to sequencing depth and detection thresholds. Meteor consistently produced slightly lower FAVA values than sylph, reflecting the greater stability conferred by its curated saliva-specific reference compared with sylph’s broader database, which captures more low-abundance taxa. This difference in within-group variability between classifiers mirrors the beta-diversity and richness results: sylph’s sensitivity to rare, depth-dependent detections introduces variability that persists across normalization strategies, reinforcing that classifier choice is an independent source of analytical variance in cross-cohort microbiome studies.

**Figure 4.**
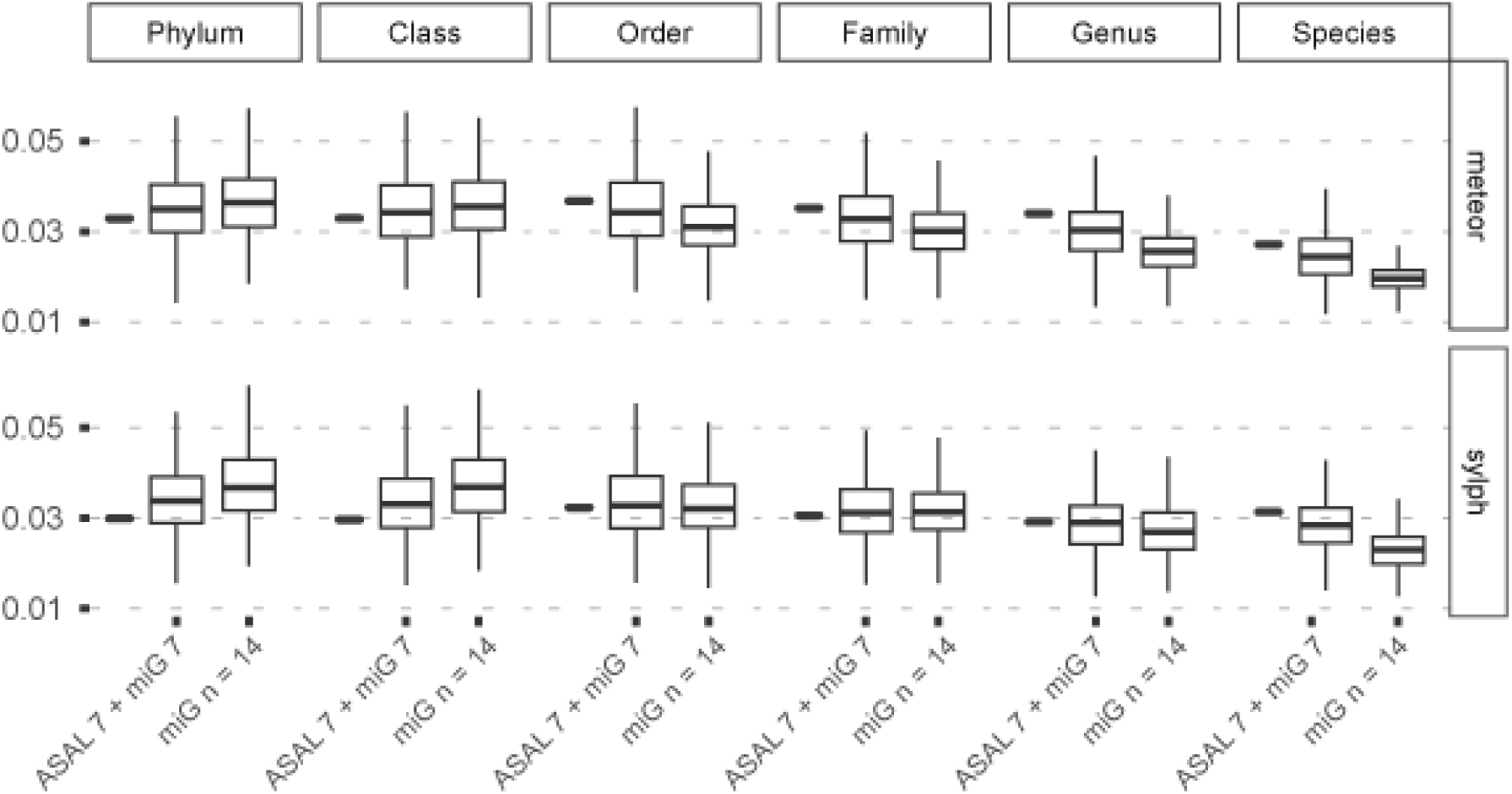
FAVA-based assessment of salivary microbiome variability across taxonomic ranks and profiling methods using rarefied profiles. Boxplots show the distribution of FAVA scores across 500 bootstrap replicates for two configurations: miG (n = 14) and the Mixed (7 ASAL + 7 miG) group. Results are shown for each taxonomic level (phylum to species). At the beginning of each panel, short horizontal bars represent the single, non-bootstrapped FAVA values computed for the ASAL dataset (n = 14) using each method. Higher FAVA values indicate greater inter-individual compositional variability, while lower values reflect more stable microbial community structures within groups.

Rarefied datasets showed uniformly higher FAVA values than unrarefied ones (Supplementary Figure 1), in line with the stochastic loss of rare taxa during downsampling. As previously reported(29), reduced microbial richness, whether due to technical factors such as rarefaction or biological perturbations, correlates with increased inter-individual variability, a trend similarly reflected in the FAVA patterns observed here.

### Coverage-based assessment of challenging taxa and extraction biases

Finally, we evaluated whether the extraction protocol affected the detection of structurally or physiologically challenging taxa by comparing genome coverage distributions between ASAL and miG samples. We focused on meteor-derived profiles identified in prior analyses as the most controlled and directly comparable across protocols. Out of 592 metagenomic species pangenomes (MSPs), 12 showed statistically significant differences (adjusted p < 0.05 in Kolmogorov–Smirnov and/or Cramér–von Mises tests; Table 1, Figure 5). A subset of the significant MSPs identified had low mean coverage or prevalence. In these cases, the distributional shift was primarily driven by detection in miG and a near-complete absence in ASAL (e.g., Selenomonadaceae sp. 905372875). Consistent with previous results, this asymmetry reflects the deeper sequencing depth of the miG dataset, which enables recovery of low-abundance or structurally challenging taxa that remained undetected in ASAL, despite rarefaction. These results likely reflect differences in detection sensitivity driven by sequencing effort. Saliva is known to contain a high proportion of host DNA, often exceeding 90% of total reads (14), which can further reduce microbial signal and taxonomic resolution when sequencing depth is limited (30).

**Figure 5.**
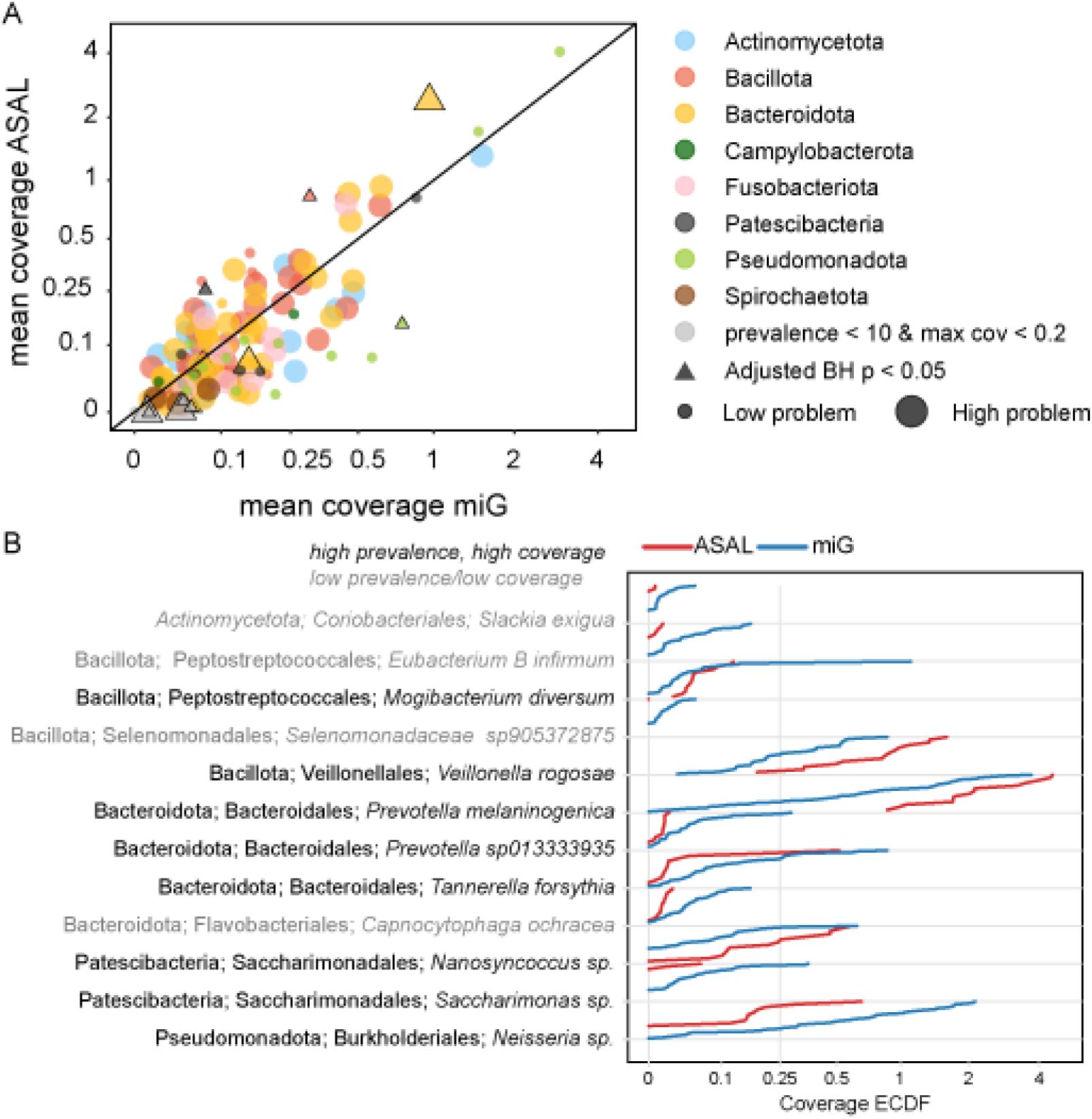
Comparison of metagenomic species pangenome (MSP) coverage across groups. (A) Mean coverage per MSP, computed using rarefied reads and meteor, across samples from miG (x-axis) and ASAL (y-axis). Points are colored by phylum and scaled by extraction protocol sensitivity, with larger circles indicating MSPs belonging to orders flagged as problematic (see Table 1). Triangles denote MSPs with significant differences between protocols (adjusted BH p < 0.05), as determined by either the Kolmogorov–Smirnov or Cramér–von Mises test. Grey triangles indicate significant MSPs with low prevalence (<10 samples) or low maximum coverage (<0.2). (B) Empirical cumulative distribution functions (ECDFs) of MSPs with significant differences between protocols. Each panel compares ECDF coverage distributions in ASAL (red) and miG (blue). Taxa names are shown in Phylum; Order; *Species* format, with greyed labels indicating low-prevalence, low-coverage MSPs. In panel B, ECDF curves are colored red (ASAL) and blue (miG) to facilitate direct protocol comparison; phylum-level coloring as in panel A is not applied to avoid visual overlap across the twelve taxa shown.

**Table 1.** Comparison of salivary microbial taxa between miG and ASAL cohorts. The table presents the prevalence (Prev) and mean coverage (Mean) of selected species-level taxa in the miG and ASAL groups. Taxa are listed by their complete taxonomy (Phylum; Order; *Species*). Statistical comparisons include the Kolmogorov–Smirnov test (*KS_p*) and centered log-ratio-transformed Mann–Whitney test (*CM_p*).

| Taxonomy<br>Phylum; Order; *Species* | Prev<br>miG | Mean<br>miG | Prev<br>ASAL | Mean<br>ASAL | KS_p | CM_p |
| --- | --- | --- | --- | --- | --- | --- |
| Actinomycetota; Coriobacteriales; *Slackia exigua* | 25 | 0.011 | 2 | 0.001 | 0.021 | 0.131 |
| Bacillota; Peptostreptococcales; *Eubacterium_B infirmum* | 32 | 0.045 | 5 | 0.004 | 0.016 | 0.131 |
| Bacillota; Peptostreptococcales; *Mogibacterium diversum* | 33 | 0.073 | 14 | 0.063 | 0.041 | 0.35 |
| Bacillota; Selenomonadales; *Selenomonadaceae CAJPQU01 sp905372875* | 25 | 0.012 | 0 | 0 | 0.016 | 0 |
| Bacillota; Veillonellales; *Veillonella rogosae* | 39 | 0.307 | 14 | 0.84 | 0.025 | 0 |
| Bacteroidota; Bacteroidales; *Prevotella melaninogenica* | 38 | 0.962 | 14 | 2.443 | 0.032 | 0 |
| Bacteroidota; Bacteroidales; *Prevotella sp013333935* | 35 | 0.052 | 11 | 0.011 | 0.02 | 0.182 |
| Bacteroidota; Bacteroidales; *Tannerella forsythia* | 37 | 0.15 | 12 | 0.068 | 0.049 | 0.362 |
| Bacteroidota; Flavobacteriales; *Capnocytophaga ochracea* | 35 | 0.048 | 12 | 0.013 | 0.016 | 0 |
| Patescibacteria; Saccharimonadales; *Nanosyncoccus* sp. | 23 | 0.076 | 13 | 0.257 | 0.073 | 0 |
| Patescibacteria; Saccharimonadales; *Saccharimonas* sp. | 27 | 0.057 | 2 | 0.006 | 0.049 | 0.182 |
| Pseudomonadota; Burkholderiales; *Neisseria* sp. | 38 | 0.754 | 9 | 0.15 | 0.025 | 0.131 |

In addition to low-prevalence or low-coverage cases, taxa such as *Veillonella rogosae*, *Prevotella melaninogenica*, and *Neisseria sp.*, core members of the salivary microbiome, exhibited subtle distributional shifts, often with slightly higher or more uniform coverage in ASAL (Table 1, Figure 5). These genera are highly prevalent and abundant across healthy cohorts, collectively accounting for up to ∼30% of oral microbial reads (31), and are generally recovered reliably across protocols. The small number and limited effect size of significantly shifted MSPs, many of which were not associated with structurally resilient or hard-to-lyse taxa, suggest that deep sequencing in miG largely compensates for the absence of mechanical lysis. The remaining differences are likely driven by a combination of minor extraction biases, ecological variation, and stochastic effects near detection thresholds that influence abundance more than presence.

### Study limitations

Although this study demonstrates the potential of host-sequencing datasets for oral microbiome analysis, several limitations should be acknowledged. The ASAL and miG cohorts differed not only in extraction protocols but also in collection kits, sequencing platforms, and library preparation workflows, each of which could introduce technical bias. Inter-individual and cohort-specific factors, such as age, medication use, or environmental exposures, may also contribute to the observed differences in microbial composition. Nevertheless, these differences were limited in our study, likely reflecting the stability of the core salivary microbiome. This is illustrated by dominant and ubiquitous taxa such as *Streptococcus*, *Neisseria sp.*, and *Prevotella*, which persist across individuals and methodological variations (32). In addition, the retrospective nature of the miG dataset precluded experimental control over storage time and environmental conditions, both of which are known to affect DNA integrity and microbial representation. Finally, the relatively small size of the ASAL dataset limited statistical power for rare taxa, and the heterogeneous sequencing depths across samples constrained our ability to fully disentangle biological variation from technical noise.

### Best practices for future salivary metagenomics using host-oriented data

The recent landmark study by Kamitaki et al.(4), which profiled oral microbiomes from 12,519 individuals by reanalyzing unmapped reads from saliva-derived WGS, demonstrates the scientific potential of precisely the approach we validate here. However, their microbiome pipeline validation remained notably limited: profiles were assessed using a single principal coordinate analysis based on Bray–Curtis distances, comparing SPARK species abundances with Human Microbiome Project oral samples, confirming only that SPARK profiles broadly clustered with oral rather than gut communities. While this confirms taxonomic plausibility, it does not address whether the host-centric extraction protocol itself introduces systematic biases, nor does it evaluate compositional stability across samples, sensitivity to sequencing depth, or differential detection of structurally challenging taxa. In contrast, our study provides direct experimental evidence that such validation requires systematic comparison of host-optimized and microbiome-optimized extraction protocols across two complementary profiling methods, rarefied and unrarefied conditions, and coverage-based statistical testing of extraction-sensitive orders.

Building on these findings, several best practices can guide future efforts to profile the salivary microbiome from host-oriented or retrospective sequencing data. First, harmonizing sequencing depth is essential: the present results indicate that at least 30–40 million total reads per sample are needed for reliable detection of low-abundance taxa and stable diversity estimates. Second, platform-specific and library-preparation effects should be carefully documented and, where possible, corrected using matched controls or normalization strategies. Third, researchers should adopt consistent bioinformatic workflows, including standardized quality filtering, host read removal, and taxonomic classification using unified reference databases, to minimize methodological noise. Fourth, classifier choice should be treated as an independent analytical variable rather than an interchangeable implementation detail. Our results show that coverage-based classifiers against curated, community-specific databases (such as meteor against the oral MSP catalog) produce profiles that converge across depth-mismatched datasets after rarefaction, whereas sketch-based classifiers against broad reference databases (such as sylph against GTDB) retain systematic detection asymmetries that rarefaction does not resolve. For cross-cohort studies combining host-centric and microbiome-optimized samples, coverage-based approaches are preferable for compositional comparisons; sketch-based classifiers are better suited to discovery analyses where maximizing taxonomic recall is the priority. Fifth, the application of bootstrapping or rarefaction-based normalization provides a rigorous framework for evaluating the robustness of microbiome signals across heterogeneous datasets. Finally, comprehensive metadata reporting, encompassing sequencing depth, platform, and analytical parameters, is crucial for reproducibility and facilitating effective cross-cohort comparisons.

Adhering to these practices will improve the comparability and interpretability of salivary metagenomic studies, ensuring that the vast archives of host genomic data can be confidently leveraged to explore microbial diversity, stability, and host–microbiome interactions at the population scale.

Applied at the scale of major genomic biobanks, where saliva-based WGS data from tens of thousands of participants already exist, the approach described here could enable retrospective microbiome studies of unprecedented size. This would facilitate microbiome genome-wide association studies (mGWAS), longitudinal analyses of oral microbial diversity in relation to aging, disease, and medication use, and cross-cohort comparisons of population-level microbial variation, all without generating new samples or incurring sequencing costs beyond those already invested.

## Conclusion

The vast archives of saliva-based whole-genome sequencing data generated by population genomic biobanks represent an underutilized resource for microbiome research. This study demonstrates that the standard host-oriented extraction and sequencing workflows used in these settings can yield robust, reproducible salivary microbiome profiles — without any modification to existing protocols.

Sequencing depth, rather than extraction optimization, emerged as the primary driver of microbiome community stability, and only 2% of detected taxa showed protocol-related differences. These findings establish a proof of concept for the large-scale recovery of microbiome data from biobank whole-genome sequencing data obtained from saliva collection kits. It opens the door to population-scale host–microbiome analyses that integrate genomic and microbial data from the same individuals at no additional sampling cost.

## Methods

### Sample collection

Two anonymous datasets were compared in this study. The first dataset, referred to as miG, consisted of whole-genome sequence (WGS) data derived from saliva samples collected from 39 individuals of the GAZEL cohort (24, 33) using the OG600 saliva collection kit (DNA Genotek™). These data were generated initially for host whole-genome sequencing, and the present analysis reuses the non-human (microbial) reads for metagenomic profiling. DNA extraction was performed at the University Hospital of Brest using magnetic bead technology on a Maxwell (PROMEGA™) instrument. Shotgun metagenomic libraries were prepared using the Illumina™ TruSeq PCR-free protocol with 1 µg of input DNA and sequenced on an Illumina™ NovaSeq 6000 platform (for more details, see (24)). Raw reads were processed using BBDuk (v.36.02) for adapter trimming and quality filtering, and Kraken2 (v.2.0.7) with the GRCh38 human reference genome used to remove human reads before microbial analysis.

The second dataset, hereafter referred to as ASAL, was obtained from the public repository of the study *“Evaluation of an Adapted Semi-Automated DNA Extraction for Human Salivary Shotgun Metagenomics”* (34). Saliva samples were collected using OMNIgene•ORAL OM-501 kits (DNA Genotek™), and DNA was extracted using two distinct protocols: a semi-automated method (Protocol 1: MGP SOP 01, adapted from the International Human Microbiome Standards, https://mgps.eu/sops/mgp-sop-001-v1/) and a manual method (Protocol 2: QIAGEN QIAamp PowerFecal Pro kit). A total of 14 samples were available and included in this analysis. Of these, 8 were raw saliva samples collected from individual donors, and 6 were pooled samples, each derived by combining saliva from two individuals. To test both extraction protocols, pooled samples were split into two aliquots: one processed using the semi-automated MGP SOP 01 V1 protocol and the other using the manual QIAGEN QIAamp PowerFecal Pro kit. All ASAL samples were sequenced on the Ion Proton platform and made publicly available through the European Nucleotide Archive (ENA) under accession PRJEB57658. Processed metagenomic reads from these samples, with human sequences already removed, were used in our comparison.

### Bioinformatics pipeline

The complete bioinformatics pipeline, including both classifiers and normalization strategies, is illustrated in Figure 1. Both datasets were analyzed using the meteor (v2.0.18) pipeline (https://github.com/metagenopolis/meteor), a comprehensive tool for metagenomic taxonomic profiling that assigns reads through direct mapping to reference genomes and infers taxonomic composition from genome-wide coverage patterns. The pipeline was run with its saliva-specific microbial reference database (https://zenodo.org/records/14181351), which includes a curated set of 853 metagenomic species pangenomes (MSPs) tailored for high-confidence detection of salivary taxa. While this targeted reference enables consistent profiling of the salivary core microbiota and provides high-confidence species-level assignments when the corresponding genomes are represented, it may underrepresent rare or novel taxa absent from the database. Reads were imported using ‘meteor fastq’ with paired-end mode. Mapping was performed against the oral reference catalog (hs_8_4_oral) using end-to-end alignment (bowtie2), a minimum nucleotide identity threshold of 0.95, a maximum of 10,000 alignments per read, read trimming at 80 bp, and a smart-shared read counting strategy (‘meteor mapping –c smart_shared –t 6’). Species abundance profiles were computed using coverage normalization, a minimum MSP core gene detection threshold of 10% (––msp_filter 0.1), and 100 core genes per MSP (––core_size 100), with no rarefaction applied at the profiling stage (‘meteor profile –l –1’). Samples were subsequently merged using ‘meteor merge’.

To complement this approach, we also performed taxonomic classification using sylph (v0.8.0) and sylph-tax (v1.5.0) (https://github.com/bluenote-1577/sylph; https://github.com/bluenote-1577/sylph-tax). Sylph employs a minimizer-based mapping strategy to assign reads to the Genome Taxonomy Database (GTDB; release R220, April 2024), enabling sensitive detection across a broad phylogenetic spectrum and improving the recovery of uncharacterized salivary taxa. Profiling was performed by running sylph profile against the GTDB-R220 pre-built database (gtdb-r220-c200-dbv1.syldb) with 20 threads, passing paired-end reads for the miG dataset (-1 *R1.fastq.gz –2 *R2.fastq.gz) and single-end reads for the ASAL dataset. Taxonomic annotation was subsequently performed using sylph-tax (sylph-tax taxprof *.tsv –t GTDB_r220 –-add-folder-information), and per-sample profiles were merged into a single abundance table (sylph-tax merge –-output comparison.tsv –-column relative_abundance).

To assess the influence of sequencing depth and normalization methods, two strategies were applied: (1) rarefaction of both datasets to 10⁶ reads, and (2) no rarefaction. The threshold of 10⁶ reads was selected to retain all samples from both datasets (minimum observed depth: 1.5 million reads in the ASAL dataset) while substantially reducing the sequencing depth advantage of the miG dataset and enabling community composition comparison under matched sampling effort. This threshold was consistent with prior salivary metagenomics studies and sufficient to recover the dominant core microbiota in both profiling methods.

## Statistical Analysis

Species richness, defined as the number of taxa detected per sample, was computed independently for each taxonomic profiling method (meteor and sylph), using both rarefied and unrarefied datasets. Differences in richness between the ASAL and miG groups were tested using Wilcoxon rank-sum tests. Taxon-level overlap across datasets was visualized using Venn diagrams to compare detection breadth between groups and profiling methods.

Microbial community structure was examined using pairwise beta-diversity distances calculated with the Jaccard distance, which reflects presence–absence variation. Principal Coordinates Analysis (PCoA) was applied to each distance matrix using classical multidimensional scaling. Group-level differences between ASAL and miG samples were evaluated using permutational multivariate analysis of variance (PERMANOVA), and within-group dispersion was assessed via multivariate dispersion analysis (BETADISPER), followed by ANOVA. All richness and beta-diversity comparisons were performed on both rarefied and unrarefied data to evaluate the impact of sequencing-depth normalization.

In addition to pairwise metrics, we applied FAVA (F-statistic-based Analysis of Variability in Abundances) to quantify compositional variability within each group (29). Unlike beta-diversity approaches that assess differences between individual sample pairs, FAVA estimates the degree of inter-individual variation across all samples within a group. Analyses were performed using relative abundance profiles spanning taxonomic levels from phylum to species level. To enable balanced comparisons between groups of unequal size, we downsampled the larger miG group (n = 39) to match the ASAL group size (n = 14) across 500 bootstrap replicates (set.seed(42)), yielding bootstrapped estimates of FAVA scores in the miG group. In addition, we generated a mixed bootstrap group comprising 7 ASAL and 7 miG samples by resampling 500 times. This design allowed us to assess the influence of cross-cohort mixing on within-group compositional stability while controlling for group size and sequencing depth.

To assess whether differences in extraction protocols influenced the detection of structurally or physiologically challenging taxa, we focused on a subset of bacterial Orders known to be resistant to standard lysis methods (Supplementary Table 1). For each taxon, we compared the distribution of genome-coverage values obtained with the meteor pipeline between miG and ASAL rarefied samples. These coverage-based profiles reflect quantitative detection sensitivity and genome representation. Distributions were compared using the Kolmogorov–Smirnov (K–S) and Cramér–von Mises tests. To account for multiple comparisons, p-values were adjusted using the Benjamini–Hochberg across all 1,184 tests (592 MSPs × 2 statistical tests). This analysis allowed us to assess whether one of the protocols favored the detection of specific taxa beyond the overall community-level patterns captured by diversity and composition metrics.

All statistical analyses and visualizations were performed using R (v 4.3.1). Data wrangling and visualization relied on the tidyverse suite, with additional functions from the janitor (https://sfirke.github.io/janitor/index.html), stringr (https://stringr.tidyverse.org/), and ggpubr (https://rpkgs.datanovia.com/ggpubr/) packages, as well as base R. Compositional variability was assessed using the FAVA R Package v.1.0.0. All analysis scripts and statistical workflows are openly available in a dedicated GitHub repository (https://github.com/lvelosuarez/miG-notebook.git) and have been archived on Zenodo (https://zenodo.org/records/15881253).

## Data availability

The taxonomy tables supporting this study are openly available in the dedicated GitHub repository (https://github.com/lvelosuarez/miG-notebook.git) and archived on Zenodo (https://zenodo.org/records/15881253). The ASAL dataset is publicly available through the European Nucleotide Archive (ENA) under accession PRJEB57658. Raw sequencing reads from the miG (GAZEL cohort) dataset cannot be shared publicly due to restrictions imposed by the French data protection framework (RGPD/GDPR) and participant consent terms applicable to the GAZEL-ADN study; access may be granted upon reasonable request, subject to a data access agreement with the study principal investigators.

## Acknowledgements

We thank all members of the GAZEL cohort for their participation in the GAZEL-ADN study and for enabling the generation of the miG dataset. This work received financial support from the French Ministry of Research and Innovation through the POPGEN project of the French Medical Genomics Plan. We are also grateful to the Clinical Investigation Center (CIC) of CHU Brest for DNA sample collection and extraction, and to the CNRGH-CEA for performing the whole-genome sequencing.

## Author contributions

Lourdes Velo-Suárez (L.V.-S.)

Conceptualization; Methodology; Formal analysis; Data curation; Visualization; Writing – original draft; Writing – review & editing.

Anthony F. Herzig (A.F.H.)

Formal analysis; Validation; Writing – review & editing.

Ozvan Bocher (O.B.)

Formal analysis; Validation; Writing – review & editing.

Gaëlle Le Folgoc (G.L.F.)

Resources; Validation; Writing – review & editing.

Liana Le Roux (L.L.R.)

Resources, Methodology

Christelle Delmas (C. D.)

Conceptualization, Project Administration

Marie Zins (M.Z.)

Resources, Funding acquisition, Project Administration

Jean-François Deleuze (J.-F. D.)

Resources, Funding acquisition, Supervision

Geneviève Héry-Arnaud (G.H.-A.)

Writing – review & editing.

Emmanuelle Génin (E.G.)

Conceptualization; Funding acquisition; Supervision; Project administration; Writing – review & editing.

## Supplementary

**Figure 1.**
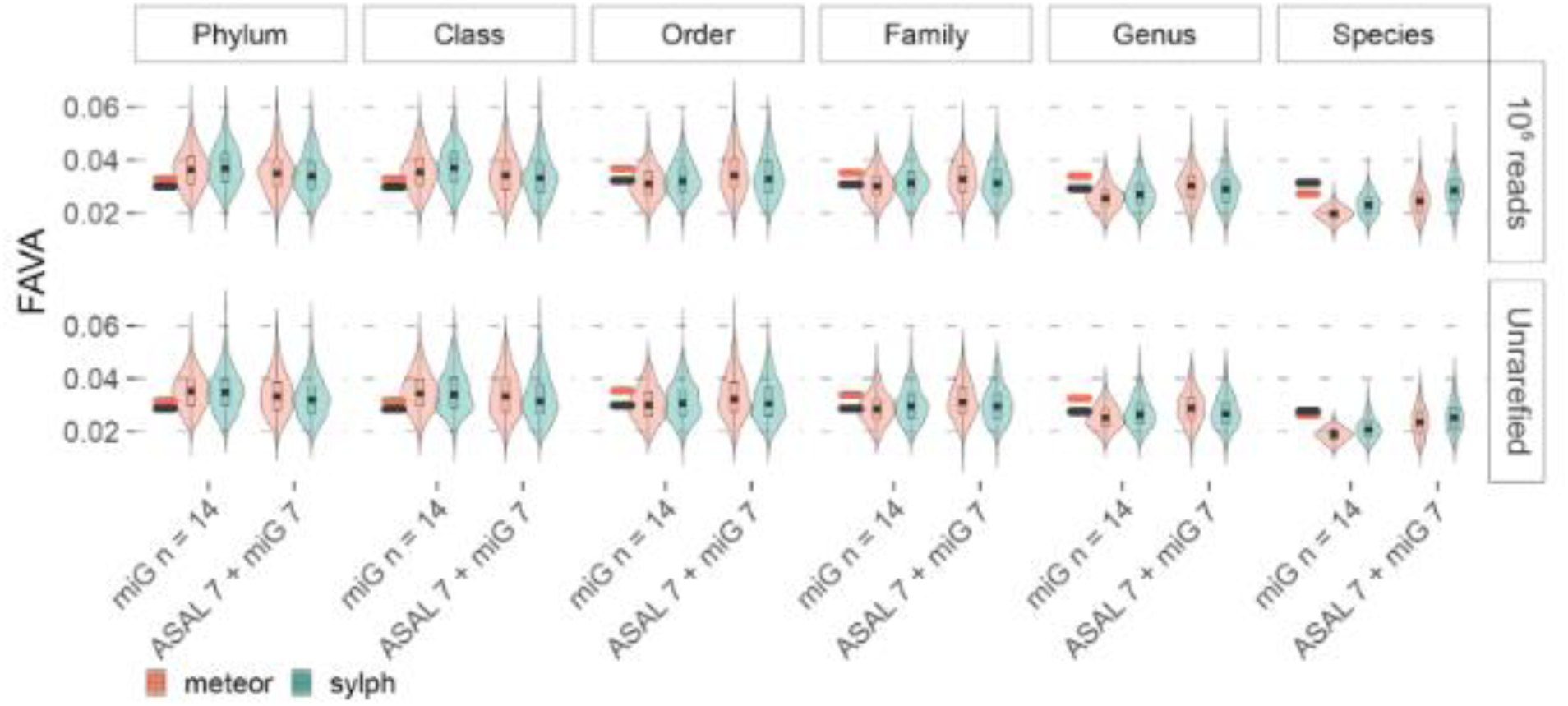
FAVA-based assessment of salivary microbiome variability across taxonomic ranks using rarefied and unrarefied profiles. Violin plots show the distribution of FAVA scores across 500 bootstrap replicates for two sample configurations: miG (n = 14) and a Mixed group composed of 7 ASAL + 7 miG samples. Results are presented for each taxonomic rank from phylum to species and for both profiling methods (meteor and sylph). Short horizontal bars at the beginning of each panel indicate the single, non-bootstrapped FAVA values computed for the full ASAL dataset (n = 14) using each method. Rarefied data (10⁶ reads) are shown in the top row, and unrarefied data in the bottom row. Higher FAVA values reflect greater inter-individual compositional variability, whereas lower values indicate more stable microbial community structures within groups.

**Supplementary Table 1.**
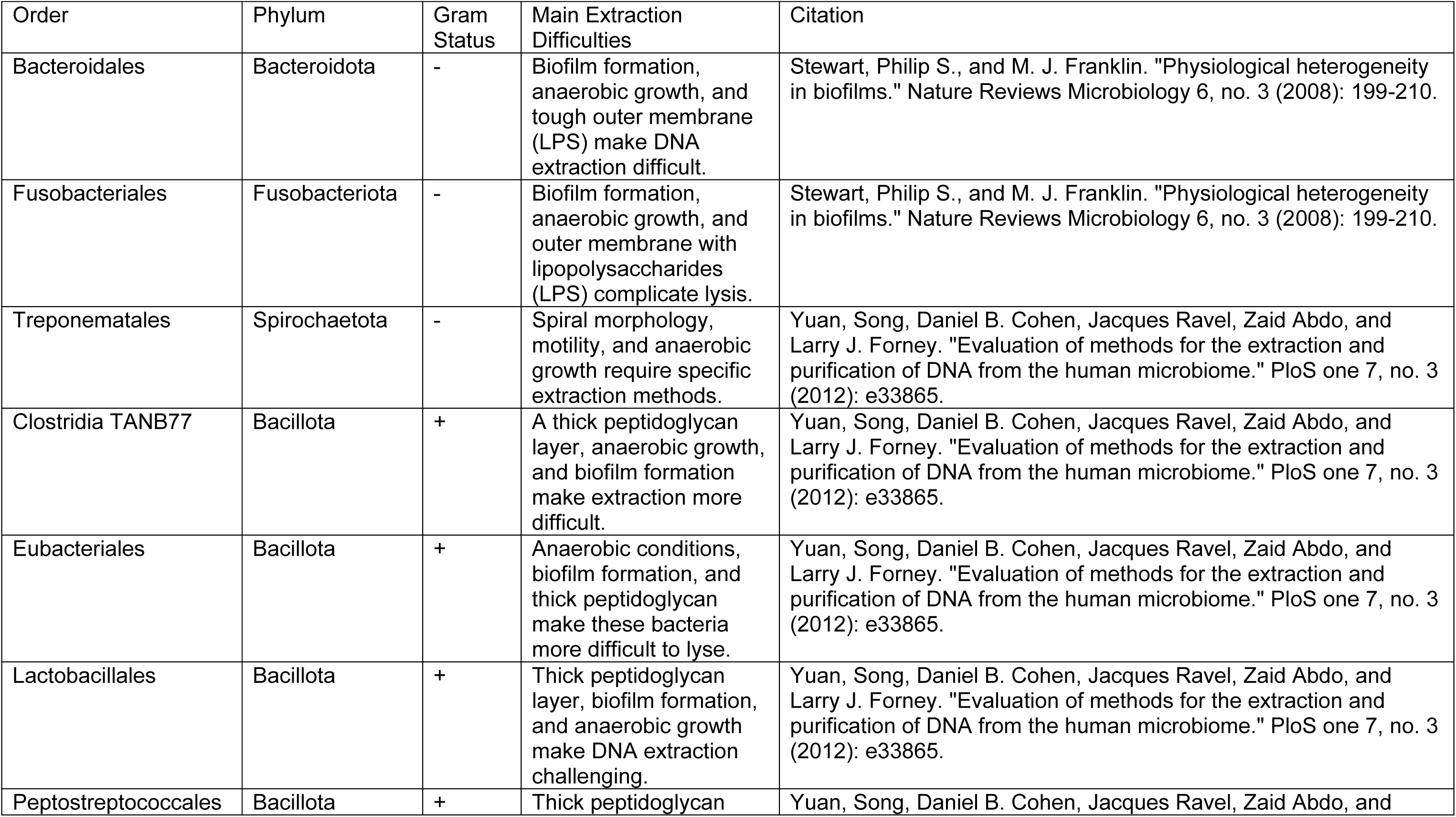
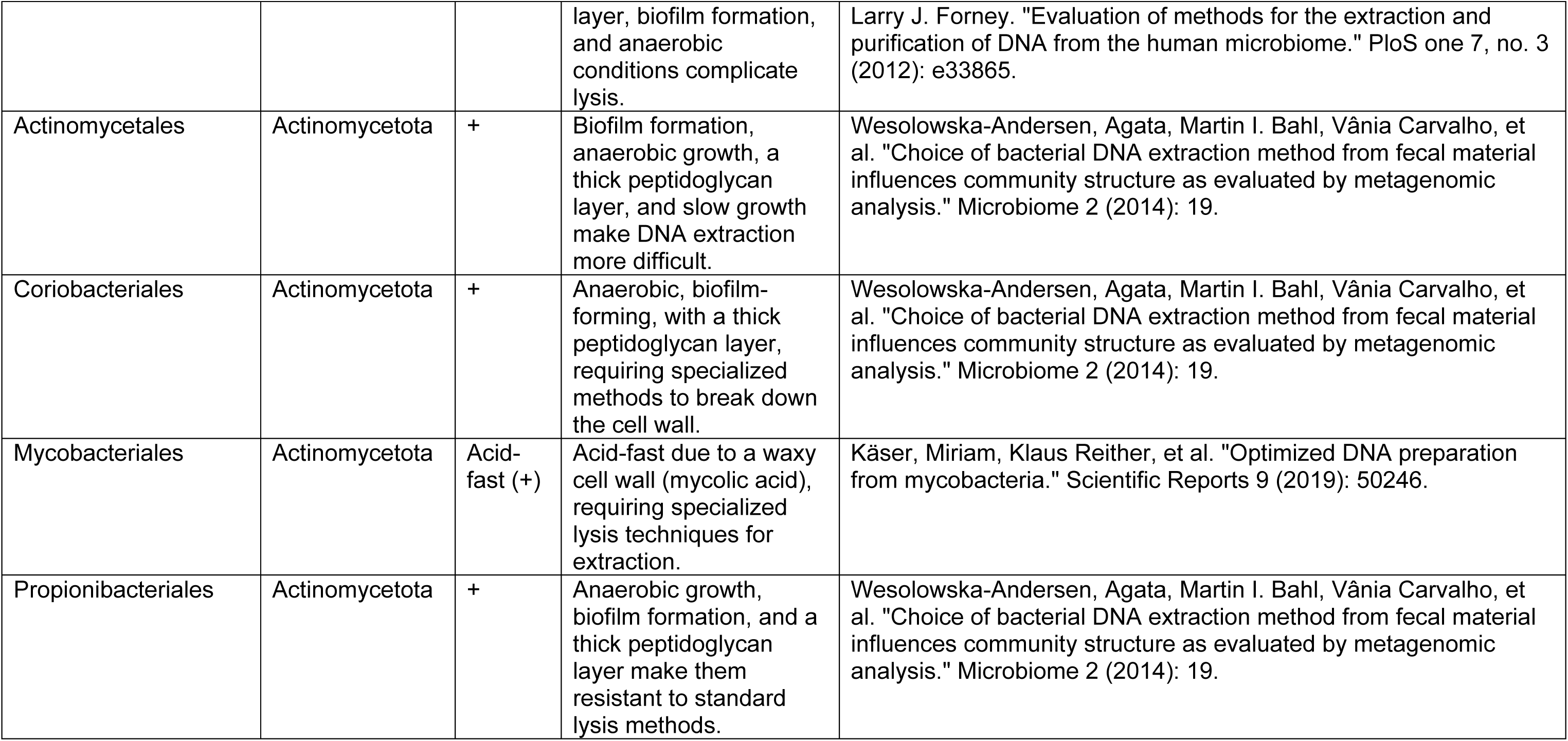
Taxonomic Orders and DNA Extraction Challenges.

| Order | Phylum | Gram Status | Main Extraction Difficulties | Citation |
| --- | --- | --- | --- | --- |
| Bacteroidales | Bacteroidota | - | Biofilm formation, anaerobic growth, and tough outer membrane (LPS) make DNA extraction difficult. | Stewart, Philip S., and M. J. Franklin. "Physiological heterogeneity in biofilms." *Nature Reviews Microbiology* 6, no. 3 (2008): 199-210. |
| Fusobacteriales | Fusobacteriota | - | Biofilm formation, anaerobic growth, and outer membrane with lipopolysaccharides (LPS) complicate lysis. | Stewart, Philip S., and M. J. Franklin. "Physiological heterogeneity in biofilms." *Nature Reviews Microbiology* 6, no. 3 (2008): 199-210. |
| Treponematales | Spirochaetota | - | Spiral morphology, motility, and anaerobic growth require specific extraction methods. | Yuan, Song, Daniel B. Cohen, Jacques Ravel, Zaid Abdo, and Larry J. Forney. "Evaluation of methods for the extraction and purification of DNA from the human microbiome." *PloS one* 7, no. 3 (2012): e33865. |
| Clostridia TANB77 | Bacillota | + | A thick peptidoglycan layer, anaerobic growth, and biofilm formation make extraction more difficult. | Yuan, Song, Daniel B. Cohen, Jacques Ravel, Zaid Abdo, and Larry J. Forney. "Evaluation of methods for the extraction and purification of DNA from the human microbiome." *PloS one* 7, no. 3 (2012): e33865. |
| Eubacteriales | Bacillota | + | Anaerobic conditions, biofilm formation, and thick peptidoglycan make these bacteria more difficult to lyse. | Yuan, Song, Daniel B. Cohen, Jacques Ravel, Zaid Abdo, and Larry J. Forney. "Evaluation of methods for the extraction and purification of DNA from the human microbiome." *PloS one* 7, no. 3 (2012): e33865. |
| Lactobacillales | Bacillota | + | Thick peptidoglycan layer, biofilm formation, and anaerobic growth make DNA extraction challenging. | Yuan, Song, Daniel B. Cohen, Jacques Ravel, Zaid Abdo, and Larry J. Forney. "Evaluation of methods for the extraction and purification of DNA from the human microbiome." *PloS one* 7, no. 3 (2012): e33865. |
| Peptostreptococcales | Bacillota | + | Thick peptidoglycan | Yuan, Song, Daniel B. Cohen, Jacques Ravel, Zaid Abdo, and |
|  |  |  | layer, biofilm formation, and anaerobic conditions complicate lysis. | Larry J. Forney. "Evaluation of methods for the extraction and purification of DNA from the human microbiome." PloS one 7, no. 3 (2012): e33865. |
| Actinomycetales | Actinomycetota | + | Biofilm formation, anaerobic growth, a thick peptidoglycan layer, and slow growth make DNA extraction more difficult. | Wesolowska-Andersen, Agata, Martin I. Bahl, Vânia Carvalho, et al. "Choice of bacterial DNA extraction method from fecal material influences community structure as evaluated by metagenomic analysis." Microbiome 2 (2014): 19. |
| Coriobacteriales | Actinomycetota | + | Anaerobic, biofilm-forming, with a thick peptidoglycan layer, requiring specialized methods to break down the cell wall. | Wesolowska-Andersen, Agata, Martin I. Bahl, Vânia Carvalho, et al. "Choice of bacterial DNA extraction method from fecal material influences community structure as evaluated by metagenomic analysis." Microbiome 2 (2014): 19. |
| Mycobacteriales | Actinomycetota | Acid-fast (+) | Acid-fast due to a waxy cell wall (mycolic acid), requiring specialized lysis techniques for extraction. | Käser, Miriam, Klaus Reither, et al. "Optimized DNA preparation from mycobacteria." Scientific Reports 9 (2019): 50246. |
| Propionibacteriales | Actinomycetota | + | Anaerobic growth, biofilm formation, and a thick peptidoglycan layer make them resistant to standard lysis methods. | Wesolowska-Andersen, Agata, Martin I. Bahl, Vânia Carvalho, et al. "Choice of bacterial DNA extraction method from fecal material influences community structure as evaluated by metagenomic analysis." Microbiome 2 (2014): 19. |

